# Remote effect of Insecticide-treated nets and the personal protection against malaria mosquito bites

**DOI:** 10.1101/073718

**Authors:** Nicolas Moiroux, Fabrice Chandre, Jean-Marc Hougard, Vincent Corbel, Cédric Pennetier

## Abstract

Experimental huts are part of the WHO process for testing and evaluation of Insecticide Treated Nets (ITN) in semi-field conditions. Experimental Hut Trials (EHTs) mostly focus on two main indicators (i.e. mortality and blood feeding reduction) that serve as efficacy criteria to obtain WHO interim recommendation. However, several other outputs that rely on counts of vectors collected in the huts are neglected although they can give useful information about vectors behavior and personal protection provided by ITNs. In particular, EHTs allow to measure the deterrent effect and personal protection of ITNs.

To provide a better assessment of ITNs efficacy, we performed a retrospective analysis of the deterrence and the personal protection against malaria transmission for 12 unwashed and 13 washed ITNs evaluated through EHTs conducted in West Africa.

A significant deterrent effect was shown for six of the 12 unwashed ITNs tested. When washed 20 times, only three ITNs had significant deterrent effect (Rate Ratios (RR)<1; p<0.05) and three showed an apparent “attractiveness” (RR>1; p<0.01). When compared to the untreated net, all unwashed ITNs showed lower number of blood-fed *Anopheles* indicating a significant personal protection (RR<1, p<0.05). However, when washed 20 times, three ITNs that were found to be attractive did not significantly reduced human-vector contact (p>0.05).

Current WHO efficacy criteria do not sufficiently take into account the deterrence effect of ITNs. Moreover the deterrence variability is rarely discussed in EHT's reports. Our findings highlighted the long range effect (deterrent or attractive) of ITNs that may have significant consequences for personal/community protection against malaria transmission. Indicators measuring the deterrence should be further considered for the evaluation of ITNs.

## Background

Between 2000 and 2015, the scale-up of malaria control interventions helped to reduce malaria mortality by 60% globally, and by 66% in sub-Saharan Africa (SSA). However, malaria is still a major cause of death with 438 000 deaths (uncertainty range: 236 000 – 635 000) of which 90% occur in SSA [1]. A recent study showed that about 70% of malaria cases were averted since 2000 due to the deployment of insecticide treated net (ITN) [2] hence underlying the need to achieve wide coverage of core interventions in all transmission settings. The ownership of ITNs increased from 2% in 2000 to 56% in 2015 but is still far from the universal coverage objective of WHO [1].

According to WHO [1], National Malaria Control Programs (NMCPs) and global malaria partners should only distribute ITNs that have been recommended by the WHO Pesticide Evaluation Scheme (WHOPES). Sixteen products are currently recommended by WHOPES [3]. WHOPES evaluation scheme is a 3 steps process (1. laboratory - 2. small- and 3. large-scale field studies) undertaken to determine the efficacy and operational acceptability of ITNs [4]. The objectives of laboratory testing (phase I) are to determine the efficacy and wash-resistance of an ITN and to study the dynamics of the insecticide on the netting fibre. Candidate ITNs that meet the requirements of phase I testing should subsequently be tested in phase II studies in experimental huts, where the efficacy of ITNs against wild free-flying mosquitoes is investigated. Candidate ITNs that reach the efficacy thresholds of phase I and phase II studies receive an interim recommendation for use as Long Lasting Insecticidal Nets (LLIN) (limited to four years of duration). To get the full recommendation, the net survivorship and attrition, fabrics physical integrity and insecticidal efficacy must be monitored and must reach WHOPES criteria during 3 years under field conditions (phase III large-scale field study) [5].

Experimental huts used in phase II studies allow evaluation of ITNs under controlled conditions that mirror those in which mosquitoes enter a human dwelling and face an ITN in normal use. Results from Experimental Hut Trials (EHTs) usually focus on two main indicators that are criteria for granting the WHO interim recommendation: the blood feeding inhibition (BFI, i.e. the reduction in blood-feeding rates relative to the control) and the mortality rates (proportion of dead mosquitoes). However, several other outputs that rely on counts of vectors collected in the huts are often neglected or analyzed with inappropriate statistical methods although they can provide useful information about vectors’ behavior and personal protection provided by ITNs. In particular, EHTs allow to measure the deterrent effect of ITNs. The deterrence is defined as the reduction in the number of mosquitoes entering the treated hut relative to the control hut (untreated nets) [4]. This indicator is measured because some insecticides (e.g. the pyrethroids) are expected to repel malaria vectors at distance preventing their entrance in the dwellings. It is therefore expected that the deterrence will be null or positive. Although it is true for most of EHTs, negative deterrence values (i.e. more malaria Anopheles were collected in the treated hut than in the control hut) occurred sometimes. In a recent review studying the impact of pyrethroid resistance in malaria vectors on the efficacy of ITN [6], the authors provide 55 values of deterrence from 17 articles reporting results of EHTs. Thirteen (24%) of these values (from 7 articles) were negatives. In this latter review, in the concerned articles [7–13] and in a recently published study [14], the authors did not discuss much about the cause or origin of these surprising “attractiveness” of treated huts. This phenomenon may have significant consequences on the efficacy of ITNs in term of personal protection against malaria transmission.

In EHT, the personal protection is defined as the reduction in the number of blood-fed mosquitoes in the treatment hut relative to the number of blood-fed mosquitoes in the control hut [4]. However, this outcome that is greatly driven by the deterrent effect of ITNs, is almost totally overlooked with the current Phase II efficacy indicators analysis process. Indeed, although current guidelines recommend calculating the personal protection of ITNs, no statistical guidance is provided to state on its significance. To illustrate the importance of the deterrence on the estimation of the personal protection of a LLIN product, we address the relationships among the BFI, the deterrence and the personal protection (see Fig 1): for a given value of BFI, the personal protection provided by an ITN could be either positive, null, or negative depending on the deterrence (see Methods for details on the mathematical relationship between the BFI, the deterrence and the personal protection).

**Fig. 1.**
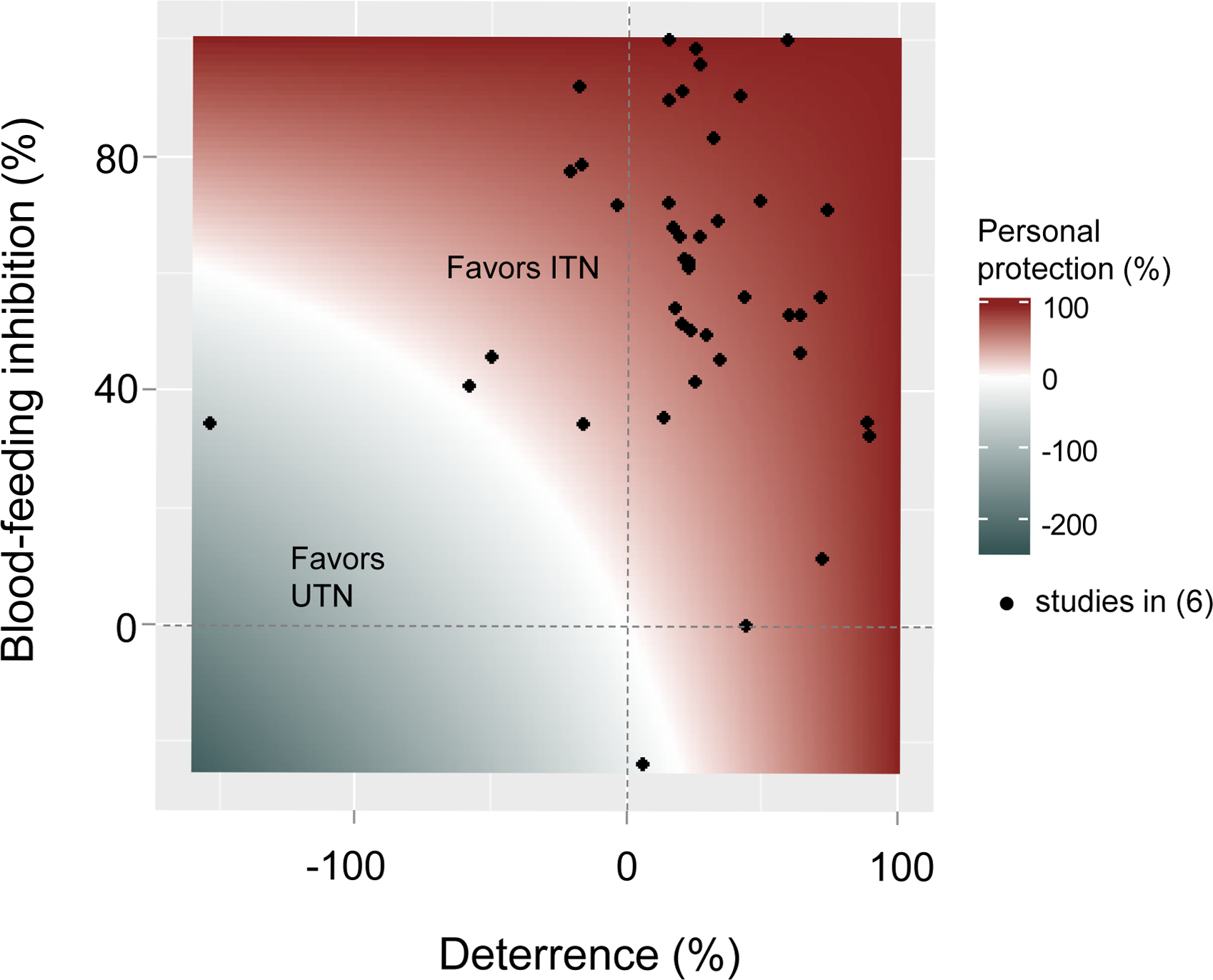
Relationships among the blood-feeding inhibition (BFI), the deterrence and the personal protection as measured in experimental hut trials. See Methods for details on the mathematical relationship among BFI, deterrence and personal protection. Values of BFI and deterrence from studies cited in the review by Strode *et al.* [6] have been plotted when both indicators can be extracted from Figure 2 and Table 12 of this review article.

Renewed interest for malaria eradication has placed greater emphasis on the development of new tools to target residual transmission (transmission that escape the control by conventional tools such as ITNs and IRS) and mosquito behavioral study are now in the spotlight [15–17]. The study of the remote effect (deterrence) and the personal protection confers by ITNs is of great importance as it might help identify weaknesses of ITNs that should be targeted by complementary vector control tools.

Therefore to provide a better assessment of ITNs used for malaria control, we performed a retrospective analysis of the deterrence and the personal protection against malaria transmission for 13 ITNs evaluated through EHTs. Trials were conducted in West Africa by Institut de Recherche pour le Développement (IRD) in the framework of the West African Anopheles, Biology and Control (ABC) network for testing and evaluation of pesticide products.

## Methods

### Calculation of the deterrence, the blood-feeding inhibition, and the personal protection (used to draw Fig 1)

The deterrence (D) is the reduction in hut entry relative to control huts (untreated nets):

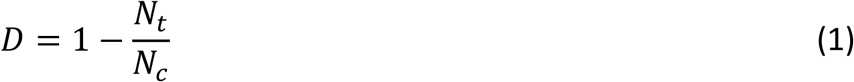

*With N_t_ the total number of mosquitoes collected in the treatment hut and veranda/exit traps and N_c_ the total number of mosquitoes in the control hut and veranda/exit traps*.

The blood-feeding inhibition (BFI) is defined as *“the reduction in blood-feeding in comparison with the control huts”* [4]. Although it is not very clear from this definition, “blood-feeding” must be understand as “blood-feeding rate” (i.e. the proportion of blood-fed mosquitoes in the huts) but not as absolute number of blood-fed mosquitoes collected in the huts. The formula commonly used to calculate the BFI is:

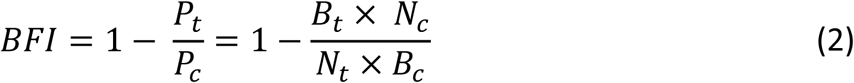

*With P_t_ the proportion of blood-fed mosquitoes in the treatment hut, P_c_ the proportion of blood-fed mosquitoes in the control hut, B_t_ the number of blood-fed mosquitoes in the treatment hut and B_c_ the number of blood-fed mosquitoes in the control hut*.

The personal protection (PP) against transmission provided by a treatment in an experimental hut study is determined by the reduction in the number of blood-fed mosquitoes in the treatment hut relative to the number of blood-fed mosquitoes in the control hut:

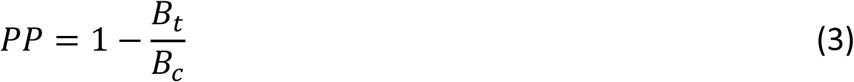

#### Relationship among PP, BFI, and D

From expression 1, we deduce expression 4:

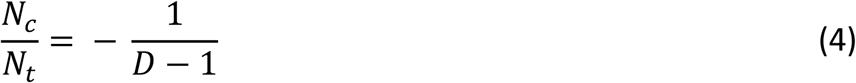

Given expression 2, we can compute BFI by solving for expression 4:

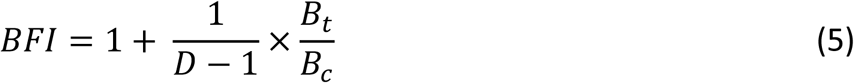

Expression 5 is equivalent to:

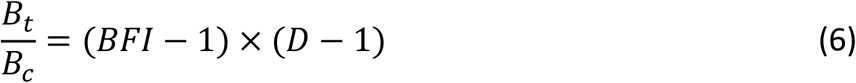

Given expression 3, we can compute PP by solving for expression 6:

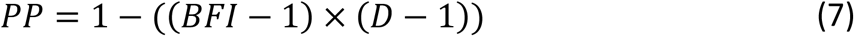

#### Studies included in the analysis

WHOPES supervised EHTs (N=10) involving 13 ITNs (12 long-lasting factory-impregnated nets and one long-lasting treatment kit (LLT) for manual impregnation) with raw data (daily collections) available for subsequent statistical analyses were included in the analysis. These studies were carried out between 2006 and 2011 in two sites (Malanville, Northern Benin and the Kou Valley, Western Burkina Faso) according to the WHO guidelines [18]. A brief description of these trials is presented in the Table 1 and a summary of the WHOPES phase II experimental hut trial protocol is provided as a supplementary material (S1 Text). Among the 13 products tested, all were tested after 20 washes and 12 were tested unwashed [19–24].

The malaria vector population in the Malanville site (North of Benin) was composed at 95 % by *An. coluzzii* (former M form) with a Kdr frequency (L1014F target-site mutation) that increased from 16 % in 2008 to 50 % in 2010 [25]. WHO cone bioassays indicated 85% and 93% mortality in 2008 [25] to the deltamethrin and permethrin insecticides, respectively. In 2010, mortality to deltamethrin decreased to 40 % [25]. In the Kou Valley (North-West of Burkina Faso), the malaria vector population was composed at 85 % by *An. gambiae s.s.* (former S form) and the Kdr frequency was 90% and the mortality rate induced by deltamethrin was 23 % [8].

#### Statistical analysis

In order to assess the deterrence, we analyzed the daily numbers of malaria vectors entering the huts using a negative binomial mixed-effect model with all the treatment arms from the 10 EHTs and the study site (Malanville or Kou Valley) as fixed effects and with the trial and the day in the trial (to deal with daily variations of the mosquito density) as nested random effects (random intercepts). The model was written as follow:

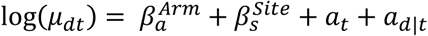

With *µ_dt_* the number of anopheles entered a particular hut on day *d* of trial *t*. 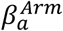 is the effect on log(*µ_dt_*) of classification in category *a* of the treatment arm and 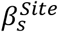 the effect of classification in category *s* (Malanville or Kou Valley) of the trial site. *a_t_* is a random intercept for trial *t* and *a_d|t_* the random intercept for day *d* of trial *t*.

**Table 1:**
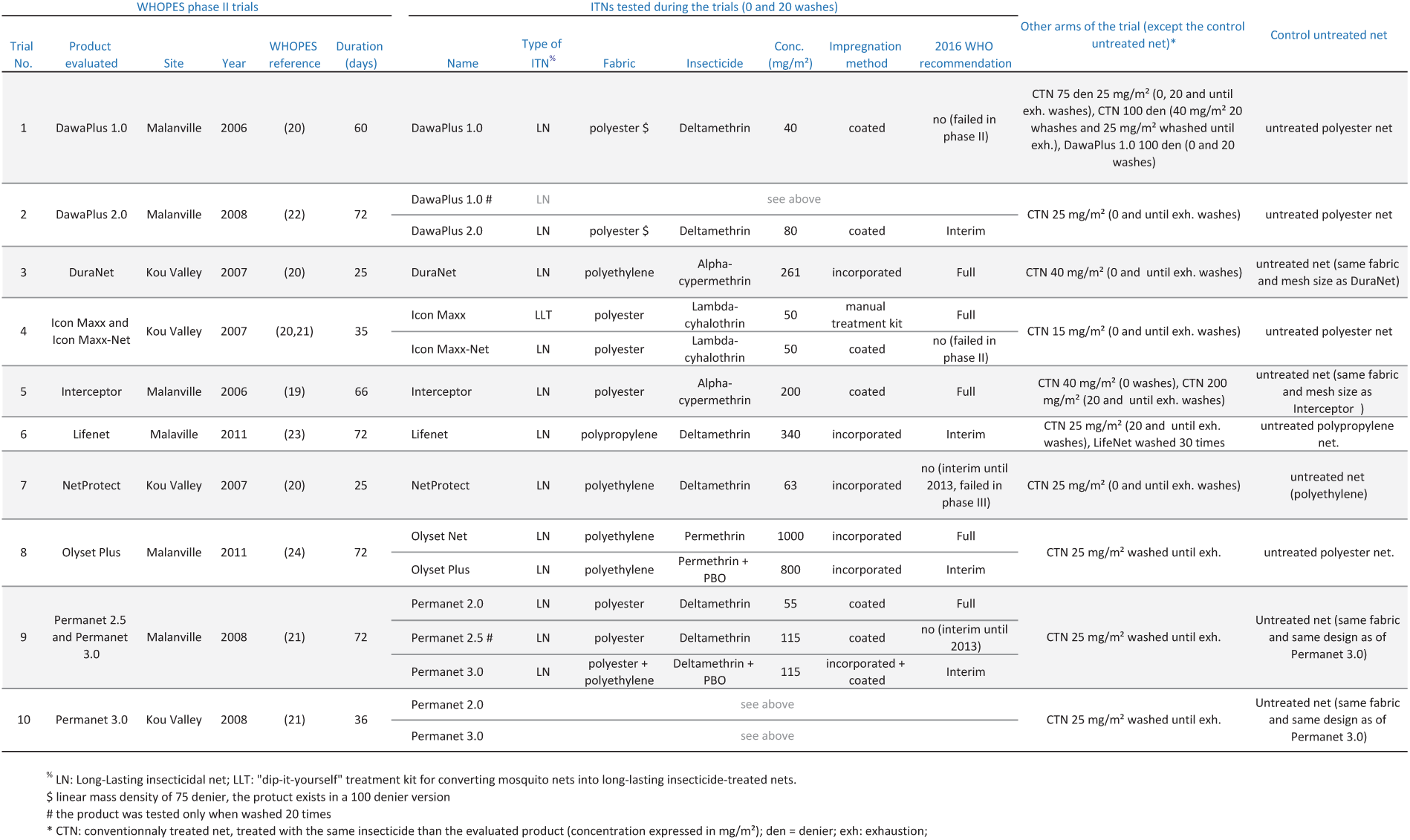
**Summary of the experimental hut trials included in the analysis**

Using the same modelling approach, we assessed the personal protection by analyzing the number of blood-fed mosquitoes collected daily in the huts. We used the ‘R’ software [26] and the additional ‘glmmADMB’ [27] package for the analysis. Rates ratios (RRs) and 95% confidence intervals were computed.

## Results

When compared to the untreated net, the number of *Anopheles* that entered the hut was lower for six of the 12 unwashed ITNs indicating a significant deterrent effect against malaria vectors (Fig 2A). For the 6 other ITNs, we were not able to detect any difference in the number of mosquito collected when compared to an UTN. When washed 20 times, only three ITNs (Interceptor, Permanet 2.0 and Permanet 3.0) had significant deterrent effect (RRs < 1; p < 0.05; Fig 2B) and three others (Icon Maxx LLT, RR= 1.59 [1.15 - 2.19], p=0.0048; Icon Maxx-Net, RR = 1.57 [1.14 - 2.16], p=0.0059; and OlysetNet, RR= 1.7 [1.18 - 2.47], p=0.0046) showed an apparent “attractiveness”. The 7 remaining ITNs did not show any difference with the untreated net (p < 0.05).

**Fig. 2.**
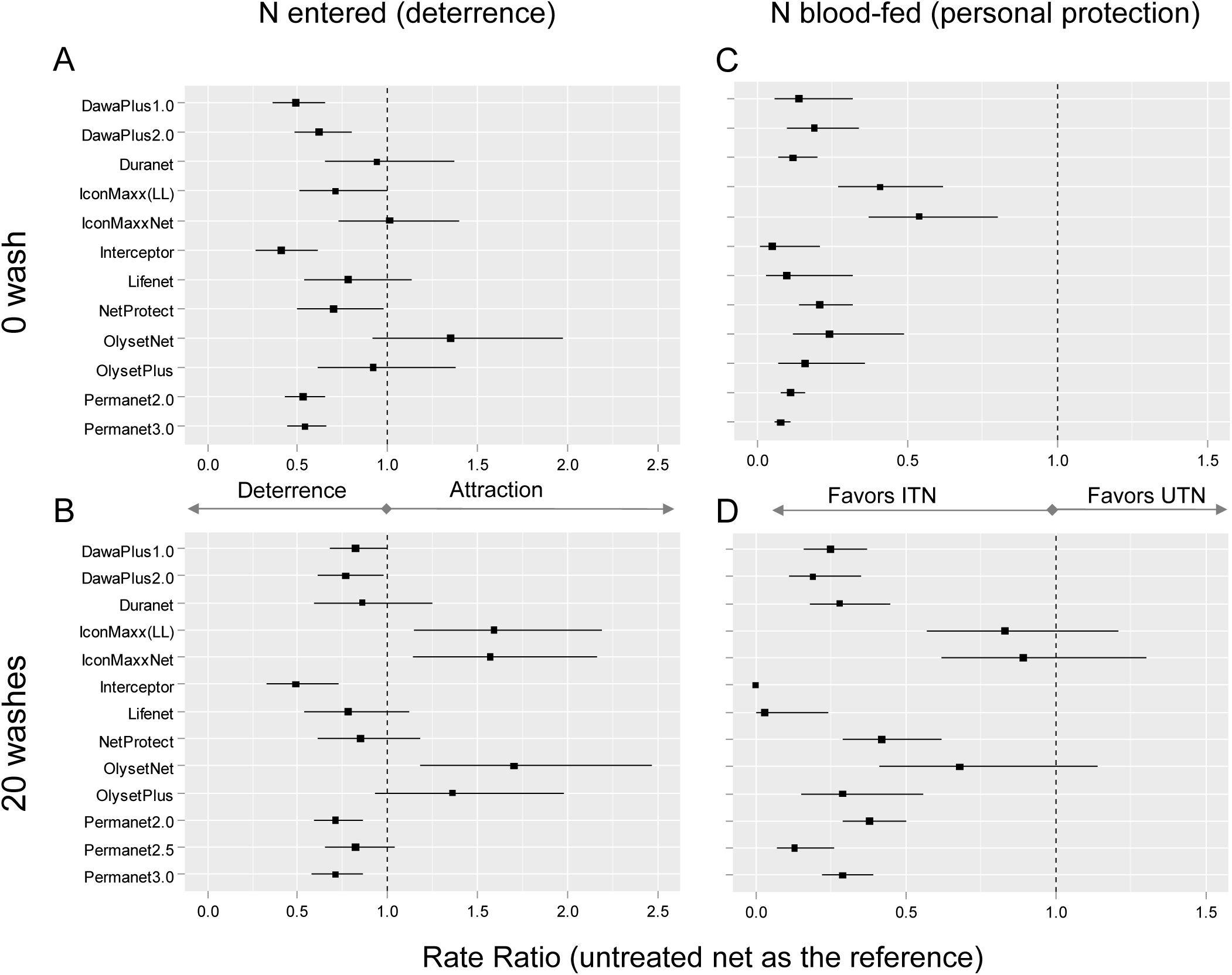
Deterrence (A,B) and personal protection (C,D) of unwashed (A,C) and washed (B,D) insecticidal treated nets evaluated through experimental hut trials in West Africa. Squares indicate Rate Ratios (with the control untreated net as reference) as obtained with Negative Binomial mixed effect models of daily counts of *Anopheles* entered the huts (A,B) and counts of blood-fed *Anopheles* (C,D). Error bars represent 95% confidence intervals of the rate ratios. LL: Long Lasting treatment for manual impregnation of the net.

When compared to the untreated net, all unwashed ITNs showed lower number of blood-fed *Anopheles* indicating a significant personal protection (RR<1, p < 0.05, Fig 2C). However when washed 20 times, the three ITNs that were found to be attractive did not significantly reduced human-vector contact when compared to an untreated net (p>0.05; Fig 2D).

## Discussion

This first analysis of the deterrence effect on personal protection of ITNs in experimental huts suggests that most, but not all of the WHO ITN recommended product tested are expected to provide personal protection against malaria transmission after 20 washes. Due to a negative deterrence effect, three ITN products did not show any significant personal protection against pyrethroid resistant malaria vectors after 20 consecutive washes. The three ITNs cause however greater killing effect on mosquito vectors than untreated nets [20,21,24].

Whatever the direction of the mosquito movement in presence of ITNs (deterrence versus attractiveness), this movement indicates that malaria vectors are able to detect the ITN at distance, before entering the hut. Deterrence of ITNs has been widely described in the last decades [28,29] because it allows reducing the vector density inside the dwellings fitted with ITN and therefore reducing the human-vector contact whether or not under the ITN. However, it is still unknown which volatiles are detected by mosquitoes. These volatiles could be the insecticide itself, additives, degradation products of these later, or products of the interaction among the insecticide, additives, CO2 and human odors. Despite a low vapor pressure (i.e. a low volatility), pyrethroids have been found in the air around a treated net [30] at concentrations (0.000021 – 0.000038 mg/m3) that are considered negligible in terms of toxicity for humans [31,32]. However, given the extraordinary sensitivity of the insects’ olfactory system [33–35], we can reasonably suspect that such concentrations might be detected by mosquitoes. This field of investigation (i.e. chemical and behavioral ecology in a context of widespread vector control tool implementation) has been neglected for decades and there is a need for more behavioral and physiological studies.

In this study, 3 ITNs of 13 that used the permethrin or the lambda-cyhalothrin insecticides were found to be attractive for malaria vectors in pyrethroid resistance areas after 20 washes. We were not able to find the same trend with corresponding unwashed ITNs indicating a significant impact of washing on ITN deterrence. The performances of an ITN can be altered by washing. After 20 washes, the mortality is strongly reduced whatever the type of ITN [19–24] indicating a reduction of the concentration of available insecticide on the net [36]. The attraction of washed ITNs might therefore indicate an insecticide dose-dependent reversal effect of orientation behavior as it has been observed for *Anopheles gambiae* with human-derived putative repellents [37] and for *Aedes albopictus* with several carboxylic acids [38,39]. Because ITNs are rarely washed 20 times in their lifetime [40–42], the kinetic of active ingredients on the fiber in relation with behavioral responses of mosquitoes are urgently needed to understand better the effect of consecutive washing on ITNs deterrence.

It should be noted that the untreated (control) net used in trial 8 (Table 1) was a polyester net, a different fabric and mesh size than the evaluated Olyset Net. To our knowledge, there is no study that address the role of net fabrics and mesh sizes of nets on human odor and CO2 dispersion. However, we cannot exclude that wide mesh ITNs (as Olyset Net [24]) allowed a better dispersion of human odor and CO2 than nets having smaller mesh size. The role of mesh size in the dispersion of odors and volatile substances would merit further investigations.

The impact of the physiological resistance to insecticide in the host-seeking behavior has been overlooked for decades. Recent findings from our team [43] showed that a lab strain of *An. gambiae* homozygous for the kdr-w mutation (L1014F) was significantly attracted by an animal host + permethrin treated net odor plume. Studies are ongoing to investigate the impact of other mutations and metabolic mechanisms conferring resistance to public health insecticides. Both the *An. gambiae* populations from Malanville and Kou Valley carried the kdr-w mutation [8,25] among other resistance mechanisms. We suspect that resistance mechanisms might modulate the host-seeking behavior by leading to the attraction of some Anopheles vectors when permethrin treated ITNs are drastically washed. Studies are ongoing to investigate the impact of other mutations and metabolic mechanisms on the behavior of mosquitoes in presence of both human host and ITNs.

We showed that three ITNs having a significant attractive effect did not provide a better personal protection than UTNs. In this particular condition, individual benefits of using these ITNs instead of an UTN (provided the UTN is maintained in good condition and is sufficiently large so that the sleeper do not make contact with it) would appeared to be null [28,44]. However, as shown in Figure 1, attractiveness would induce null or negative personal protection only for nets exhibiting a BFI rate lower than 50%.

The effect of attraction on community protection cannot be assessed precisely with EHTs data. Indeed, washed ITNs that we found to be attractive were efficient to kill an important number of mosquitoes [20,21,24] contributing to the reduction of the adult density and the lifespan of the local population of vectors. However theoretically [45], the community protection provided by an intra-domiciliary vector control tool is highly dependent on the coverage of the intervention (i.e. the proportion of people that use it) that cannot be simulated in EHTs. It is therefore impossible to conclude that the attractive property might have an effect (either positive or negative) on the community protection based on EHT outputs.

The best way to evaluate the community effect of ITNs against transmission should be to monitor and compare EIRs, malaria prevalence and incidence through a phase III Randomized Control Trial (RCT) with a negative control arm (untreated net). Nevertheless, since ITNs are now the baseline intervention for most of NMCP, the use of untreated nets as a negative control raises ethical issues. Alternatively, as compliance with ITNs is never 100%, a parasitological and clinical follow-up of non-users after the distribution of LLIN should help to measure the community effect. Community level trials are costly and time consuming and therefore the use of mathematical models of transmission using EHT data showed useful to predict community protection induced by LLIN. Such models exists and have been used to compare the potential efficacy of insecticide products having or not a repellent effect [46]. The authors of the later study found that purely toxic products with no deterrence are predicted to generally provide superior protection to non-users and even users, even if that product confers no personal protection. By extrapolation of Killeen and colleagues’ results [46], we could expect that attractive products might induce superior community protection than deterrent ones. However, according to Okumu *et al.* [47] who adapted this model to be used with EHT data, the model do not allow to deal with negative deterrence. Therefore, as a first step before community-level trials, simulations using mathematical models of transmission adapted to allow for attractive product (for example, an adaptation of the modelling approach recently published by Churcher *et al.* [48] that used EHT data to predict the impact of insecticide resistance on malaria infection) should be run to evaluate the effect of an attractive ITN at the community level. If the community effect might be confirmed, it is important to note that products which confer low or no personal protection will require adapted awareness campaign that emphasize the communal nature of protection [46].

## Conclusion

Current WHO efficacy criteria do not take into account the deterrence and the deterrence variability is neither analyzed nor discussed in the majority of the reports of experimental hut studies as illustrated in a recent literature review [6]. Consequently, there is an important gap of knowledge with unknown consequences in terms of public health. Our study points the long range effect (repellent or attractive) of ITNs, the personal protection and above all, the community protection out to be major criteria for the evaluation of ITNs.

## Acknowledgements

These Phase II trials and the present study have been run in the frame of ABC network.

## Supporting informations

**S1 Text. Summary of the WHOPES Experimental Hut Trial protocol.**

